# *3DTubularVoronoi*: a Voronoi-Based Framework for Estimating Equilibrium Cellular Packing in 3D Tubular Epithelia

**DOI:** 10.1101/2025.09.02.673513

**Authors:** Léna Guitou, Eloy T. Serrano, J. Alberto Conejero, Javier Buceta

**Affiliations:** Theoretical and Computational Systems Biology Program, Institute for Integrative Systems Biology (I2SysBio), CSIC-UV, Valencia, Spain; Instituto Universitario de Matematica Pura y Aplicada, Universitat Politècnica de València, Spain

## Abstract

Epithelial tube morphogenesis is fundamental to organ development and functioning, yet accurately modeling its intricate 3D organization and dynamics remains a challenge. In particular, current computational tools fail to capture the spontaneous emergence of non-transient scutoids, a geometric feature essential for accommodating curvature in epithelia. Here, we introduce *3DTubularVoronoi*, a novel simulation framework for modeling epithelial tube morphogenesis and determining their stable cell packing organization depending on mechanical parameters. The model employs a Voronoi tessellation strategy combined with an energy minimization approach using the Metropolis-Hasting algorithm. *3DTubularVoronoi* enables realistic epithelial tube simulations, accounting for scutoid formation and dynamic rearrangements. This tool provides an open-source, user-friendly solution for exploring gland morphogenesis and offering insights into developmental biology and tissue engineering.

**Availability:** https://github.com/TheSiMBioSysI2SysBio/3DTubularVoronoi

## I. INTRODUCTION

Epithelial tubes form the foundation of many vital organs, including the lungs, kidneys, and vasculature. As such, epithelial tube morphogenesis plays a crucial role in organ development and function. Tubulogenesis involves complex morphogenetic mechanisms influenced by mechanical forces, cell rearrangements, and tissue geometry. Gaining a deeper understanding of these intricate dynamics is essential for advancing developmental biology and unlocking new biomedical engineering applications. Given the complexity of these processes, computational modeling tools have become indispensable for systematically investigating epithelial tubulogenesis. These tools provide quantitative insights that are challenging —or even impossible at present— to achieve through experimental approaches alone.

Among the currently available 3D modeling tools, *SimuCell3D* is an efficient open-source program capable of simulating large tissues with subcellular resolution. It incorporates features such as growth, proliferation, and extracellular matrix interactions, effectively balancing resolution and computational scalability [14]. Another notable tool, *CellSim3D*, represents cells as collections of nodes connected by springs, allowing for realistic deformations and interactions, which facilitate the study of complex cellular behaviors in 3D environments [10]. Additionally, a 3D ‘bubbly’ vertex model has been employed to investigate the interplay between cell extrusion and tissue curvature, offering valuable insights into the mechanical and geometric factors shaping epithelial tissue morphology [2].

While these tools excel at capturing various aspects of epithelial tissue dynamics in 3D, none explicitly account for the spontaneous formation of *scutoids*, a geometric feature essential for efficiently accommodating curvature in epithelial tissues [5]. Existing 3D models that incorporate scutoids are either limited to flat tissue geometries [9] —restricting their applicability to curved epithelial systems—, rely on static, and purely geometrical, approaches that do not allow for the dynamic study of 3D cell packing [5, 7], or reproduce the formation of transient scutoids as cells extrude from epithelial sheets [11].

To address these limitations, we introduce a novel simulation tool based on a Voronoi tessellation modeling approach combined with a Metropolis-Hastings dynamics. Unlike previous models, our tool enables the emergence of equilibrium scutoids through a dynamic energy relaxation process and hence makes possible to find stable cell packing organization during tube formation. This advancement allows for a more accurate representation of epithelial tube morphogenesis and its cellular packing by incorporating dynamic cell rearrangements driven by mechanical forces and tissue-specific geometric constraints.

## II. MODELING APPROACH

Our modeling approach relies on a 3D solid cylinder representation with an inner cylindrical cavity simulating the body and lumen of epithelial tubes (Fig. 1A). Following previous approaches based on static Voronoi tessellations [5], our method discretizes the tubular structure into a number of sheets with constant radius, and consequently the cells, into layers along the apico-basal axis from the apical (inner) radius, *R*_*a*_, to the basal (outer) layer, *R*_*b*_ (Fig. 1B). This overcomes the computational challenges associated with 3D geodesic distances and transforms the cylindrical surfaces into their planar representations. This “unrolling” process preserves local isometry, maintaining geodesic distances between points and consequently cellular geometrical features in each radial surface such as areas and perimeters. To address the periodic boundary conditions of the cylindrical topology, we use replicas of the cylinder planar representations (see Supp. Mat.), thus avoiding issues with computing Voronoi tessellations near the edges of the unrolled cylinder. Cells in the apical surface are determined by the seeds (points) that are initially randomly distributed in the inner cylindrical surface and the Voronoi tessellation of these points determines the apical surface of cells. Subsequent cell layers are constructed by projecting normally the apical seeds towards the basal cylindrical surface. The projection length (i.e., the cell height) is calculated depending on the number of layers and the basal, *R*_*b*_, and apical, *R*_*a*_, radii (Fig.1B). In each intermediate cylindrical surface (i.e. layer) we compute the Voronoi tessellations of the seeds and each cell is defined by the set of tessellations of a given seed.

**FIG. 1.**
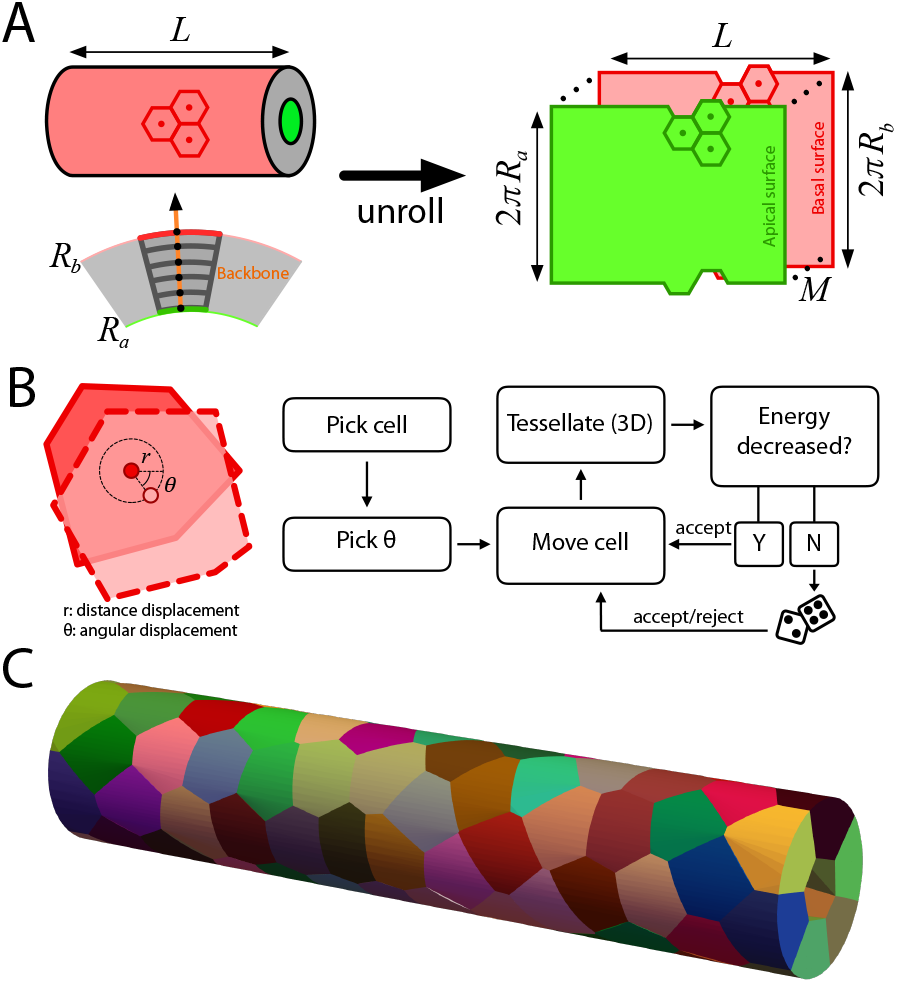
*3DTubularVoronoi*: modeling approach. **A**. Given a 3D tubular structure, the method discretizes it in a set of *M* layers, from the inner apical radius, *R*_*a*_, to the outer basal radius, *R*_*b*_, using an unrolling procedure. Each cell in the tissue is then described by the set of Voronoi tessellations in the different layers that belong to the same radial backbone. **B**. The cellular dynamics towards equilibrium packing is implemented using the Metropolis-Hasting algorithm. A random displacement of the apical seed of a cell, and therefore the displacement of all seeds defined by the cellular backbone, results in a new energy value of the tissue that is either accepted or not. **C**. 3D representation of an epithelial tube after packing equilibrium has been achieved.

Given the geometrical procedure mentioned above, we implement a dynamics based on a mechanical description as follows. We associate to each set of tessellations (i.e. cell, *C*_*i*_) an energy functional that accounts for their elastic, contractive, and adhesive components:

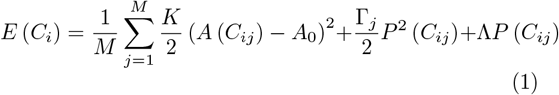

where *M* indicates the number of discretization layers, *C*_*ij*_ indicates the Voronoi region of cell *i* in layer *j, A* (*C*_*ij*_) its area (*A*_0_ being the target area), and *P* (*C*_*ij*_) its perimeter. The value of *A*_0_ can be estimated given the geometrical features of the tube where cells are embedded,

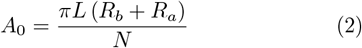

where *N* is the total number of cells in the tube and *L* its length (Supp. Mat.). In this way the energy functional follows a vertex-like description that has been shown to provide a faithful description of the mechanics of epithelial cells [3]. In this description, the parameters *K*, Γ, and Λ weigh the relative importance of elastic, contractive, and adhesive energy terms respectively. In our approach we implement an exponential decay of the contractive parameter Γ from the apical to the basal surface that can be modulated in the simulations (Supp. Mat.). Such a decrease of the contractility activity of cells from apical to basal surfaces has been recently reported by force inference approaches regardless of the cell packing configuration followed by cells [1].

Given this energetic description, we implement a relaxational dynamics using the Metropolis-Hasting algorithm (Fig.1B). Thus, choosing randomly a cell seed in the apical surface we implemented a random displacement of the apical seed of cells (see Supp. Mat. for details). After the cell seed in the apical plane is moved we compute the displacement of all the seeds defining the cell in the different layers and re-tesselate the whole tube. This leads to a new configuration of the cells in the tube and consequently a new energy of the tube, 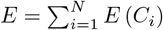, that we either accept or reject depending on the Metropolis-Hasting algorithm. Additional mathematical details are provided in the user manual (Supp. Mat.). This iterative process leads, as expected, to configurations that eventually converge to equilibrium and therefore to stable 3D cell packing (Fig. 1C).

## III. SOFTWARE IMPLEMENTATION

*3DTubularVoronoi* has been developed on the open-source R language that offers a broad accessibility and cross-platform compatibility (Windows, Mac OSX, and Linux) [12]. The code uses cluster computing to run realizations of simulation conditions in parallel, significantly reducing the computation time. Designed for accessibility, *3DTubularVoronoi* uses the user-friendly interface RStudio [13]. Parameters (dimensionless) are defined at the beginning of the code to streamline setup, and a comprehensive user manual is provided for usability (see Supp. Mat.). The tool prioritizes data generation and analysis, with inputs limited to parameter settings in the main R script and outputs provided as.RData and.csv files. Output data files are described in the manual and the code also generates plots about the energy relaxation as a function of the simulation step and as a function of the apico-basal coordinate. While not necessary for performing simulations and/or generating 3D plots of the simulated tubes, this code relies on proprietary software (a notebook developed for Wolfram Mathematica [8]) that can be run using the free, public tool Wolfram Player.

## IV. RESULTS

To illustrate the results that can be obtained from *3DTubularVoronoi*, we performed simulations inspired by the tubular epithelia found in the *Drosophila* third instar salivary glands [5]. This is a well-characterized tubulogenesis model in which cells organize around a cylindrical lumen and where the formation of scutoids has been found to be instrumental for their stability [1]. Previous approaches aimed at finding equilibrium cell packing configurations of this tissue are based on static geometrical approaches and require manual iterations of the so-called Lloyds algorithm until the results fit the experimental observations [5, 7]. Here we perform simulations using *3DTubularVoronoi*. The reported results were obtained for the following parameters: *R*_*b*_*/R*_*a*_ = 1.8, *L* = 15, *M* = 90, *N* = 8, Γ = 0.2, and Λ = 0.1.

Starting from a characteristic random seed configuration in the apical surface, we use *3DTubularVoronoi* and let the system evolve to an energy minimum. As the number of iterations (pseudo-time steps) increases, the model shows an energy relaxation (Fig. 2A), consistent with the expected behavior as cells rearrange to uniformly occupy the available tubular space. This rearrangement is mainly driven by the elastic term of the energy functional and becomes more evident when examining the average relative cell area as a function of the polygonal class in apical and basal surfaces (Fig. 2B). Thus, cells become more homogeneous in size while showing a linear increase of the average area as a function of the polygonal class as expected from the so-called Lewis law [6].

**FIG. 2.**
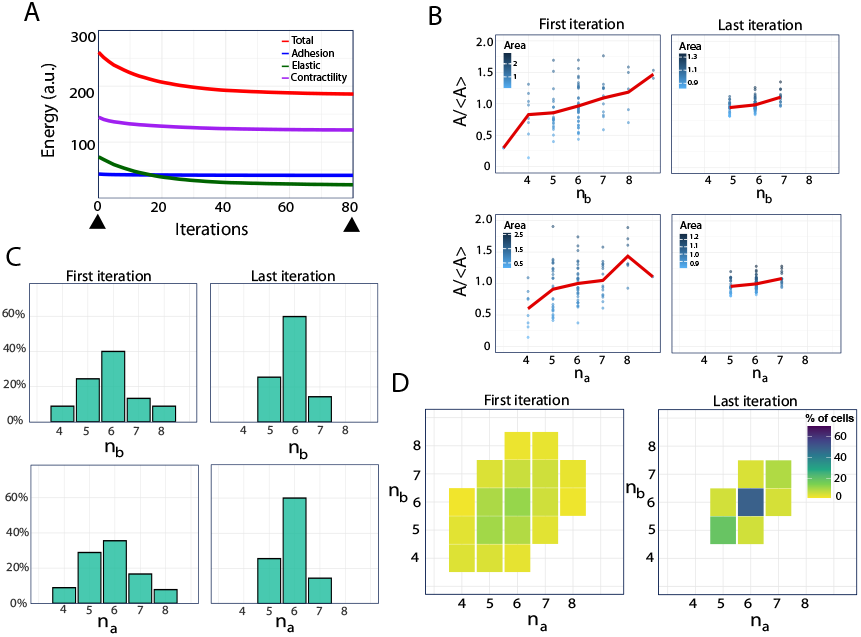
Tubular epithelia: topological properties as equilibrium approaches. **A.** Energy of the tissue as a function of the iteration number (pseudo-time). Black triangles indicate the two time points referred to as “First iteration” and “Last iteration” in panels **B**-**D** (iterations 1 and 80 respectively). **B**. Average cell area as a function of the polygonal class in the apical and basal surfaces (red solid line). Individual values are displayed by means of dots. **C**. Polygonal distributions in the apical and basal surfaces. **D**. 3D Histograms of polygonal distributions (i. e. probability of having a given number of neighbors at the apical and basal surfaces). Bin entries outside the diagonal indicate scutoids. In all panels the sample size is *n* = 10 simulations.

As for the topological arrangement/packing of cells in the tissue, we analyzed both the planar topology, in apical and basal surfaces, as well as the 3D histograms that shed light on the appearance of scutoids. As for the former, cells areas acquire topological distributions in agreement with the ones observed in the salivary glands of *Drosophila* and predicted mathematically by the Gibson-Patel distribution [4]. As for the latter, scutoids (as revealed by the out-of-diagonal entries in the histograms, see [7]) are revealed to be instrumental in the tubular architecture when energy reaches a minimum as recently shown [1]. Notably, the variability of scutoidal cells shapes, as revealed by the width of the distribution along the perpendicular direction to the diagonal, decreases as equilibrium is approached. However, the number of scutoids, as indicated by the number of cells with a different number of neighbors in the apical and basal surfaces, is consistently maintained at ∼ 75% as observed in experiments [5].

## V. CONCLUSION

In this work, we introduced *3DTubularVoronoi*, a novel computational framework for modeling epithelial tube morphogenesis and equilibrium cell packing using a Voronoi-based approach combined with Metropolis-Hastings dynamics. Our model uniquely captures the spontaneous emergence of stable scutoids, a critical geometric feature that enables epithelial tissues to accommodate curvature efficiently.

By implementing a dynamic energy minimization strategy, *3DTubularVoronoi* overcomes the limitations of previous static geometrical models and allows for a more realistic simulation of epithelial tube organization. Our results demonstrate that the model reproduces key features observed *in vivo*, including the emergence of scutoidal packing configurations and the expected energy relaxation dynamics. Furthermore, the ability to tune mechanical parameters provides a valuable tool for exploring how tissue-specific forces shape epithelial structures, offering new insights into developmental biology and tissue engineering applications.

Beyond its biological relevance, *3DTubularVoronoi* is designed as an accessible and flexible tool, implemented in R with support for parallel computing thus ensuring its usability across platforms. Future developments will focus on expanding the model capabilities to include active cellular processes (e.g. growth and division) to further enhance its predictive power for simulating tubulogenesis. In summary, *3DTubularVoronoi* provides a powerful open-source framework for researchers investigating the mechanics of epithelial morphogenesis and contributes to the broader understanding of tissue architecture in development and disease.

## FUNDING

This work was supported by the Ministerio de Ciencia e InnovaciÓn of Spain through grants PID2022-137436NB-I00 and PID2019-105566GB-I00 (J.B.). J.B. also received funding from the research network RED2022-134573-T funded by Ministerio de Ciencia e InnovaciÓn (MCIN/AEI/10.13039/501100011033) and by ‘ERDF: A way of making Europe’, by the European Union. J.B. received additional support from the E.U. COST action CA22153 ‘European Curvature and Biology Network’ (EuroCurvoBioNet). JAC is supported by Generalitat Valenciana, Project PROMETEO CIPROM/2022/21.

## ACKNOWLEDGEMENTS

We thank Adrian Nadal for fruitful comments.

## Notes

https://github.com/lenaguitou/3DTubularVoronoi

### Competing Interest Statement

The authors have declared no competing interest.

## References

[1] Anbari, S., GÓmez-Gálvez, P., Vicente-Munuera, P., Escudero, L. M., and Buceta, J. (2025). Apico-basal intercalations enable the integrity of curved epithelia. Computational and Structural Biotechnology Journal, 27, 1204–1214.

[2] Drozdowski, O. M. and Schwarz, U. S. (2024). Cell bulging and extrusion in a three-dimensional bubbly vertex model for curved epithelial sheets. arXiv preprint, 2411. 15 pages, 10 figures.

[3] Farhadifar, R., Roper, J.-C., Aigouy, B., Eaton, S., and Jülicher, F. (2007). The influence of cell mechanics, cell-cell interactions, and proliferation on epithelial packing. Current biology : CB, 17(24), 2095–2104.

[4] Gibson, M. C., Patel, A. B., Nagpal, R., and Perrimon, N. (2006). The emergence of geometric order in proliferating metazoan epithelia. Nature, 442(7106), 1038–1041.

[5] GÓmez-Gálvez, P., Vicente-Munuera, P., Tagua, A., Forja, C., Castro, A. M., Letrán, M., Valencia-ExpÓsito, A., Grima, C., Bermúdez-Gallardo, M., Serrano-Pérez-Higueras, Ó., Cavodeassi, F., Sotillos, S., Martín-Bermudo, M. D., Márquez, A., Buceta, J., and Escudero, L. M. (2018). Scutoids are a geometrical solution to three-dimensional packing of epithelia. Nature communications, 9(1), 2960.

[6] GÓmez-Gálvez, P., Vicente-Munuera, P., Anbari, S., Buceta, J., and Escudero, L. M. (2021). The complex three-dimensional organization of epithelial tissues. Development, 148(1), dev195669. REVIEW — 06 January 2021.

[7] GÓmez-Gálvez, P., Vicente-Munuera, P., Anbari, S., Marquez, A., Buceta, J., and Escudero, L. M. (2022). A quantitative biophysical principle to explain the 3d cellular connectivity in curved epithelia. Open Archive, 13, 631–643.E8. Open Archive.

[8] Inc., W. R. (2024). Mathematica, Version 14.2.

[9] Ioannou, F., Dawi, M. A., Tetley, R. J., Mao, Y., and Muñoz, J. J. (2020). Development of a new 3d hybrid model for epithelia morphogenesis. Frontiers in Bioengineering and Biotechnology, 8, 405. Original Research Article, Section: Tissue Engineering and Regenerative Medicine.

[10] Madhikar, P., Åström, J., Westerholm, J., and Karttunen, M. (2018). Cellsim3d: Gpu accelerated software for simulations of cellular growth and division in three dimensions. Computer Physics Communications, 232, 206–213.

[11] Okuda, S. and Fujimoto, K. (2020). A Mechanical Instability in Planar Epithelial Monolayers Leads to Cell Extrusion. Biophysical Journal, 118(10), 2549–2560.

[12] R Core Team (2021). R: A language and environment for statistical computing.

[13] RStudio Team (2020). Rstudio: Integrated development environment for r.

[14] Runser, S., Vetter, R., and Iber, D. (2024). Simucell3d: three-dimensional simulation of tissue mechanics with cell polarization. Nature Computational Science, 4, 299–309. Resource, Open Access.

